# Sensory modulation of gait characteristics in human locomotion: a neuromusculoskeletal modeling study

**DOI:** 10.1101/2020.12.17.423198

**Authors:** Andrea Di Russo, Dimitar Stanev, Stéphane Armand, Auke Ijspeert

**Affiliations:** Biorobotics Laboratory, École polytechnique fédérale de Lausanne, School of Engineering, Institute of Bioengineering, Lausanne, Switzerland; Willy Taillard Laboratory of Kinesiology, Geneva University Hospitals and University of Geneva, Geneva, Switzerland

## Abstract

The central nervous system of humans and animals is able to modulate the activity in the spinal cord to achieve several locomotion behaviors. Previous neuromechanical models investigated the modulation of human gait changing selected parameters belonging to the CPGs (Central Pattern Generators) feedforward oscillatory structures or to the feedback reflex circuits. CPG-based models could replicate slow and fast walking by changing only the oscillation’s properties. On the other hand, reflex-based models could achieve different behaviors mainly through optimizations of a large dimensional parameter space, but could not identify effectively individual key reflex parameters responsible for the modulation of gait characteristics. This study, investigates which reflex parameters modulate the gait characteristics through neuromechanical simulations. A recently developed reflex-based model is used to perform optimizations with different target behaviors on speed, step length and step duration in order to analyse the correlation between reflex parameters and their influence on these gait characteristics. We identified 9 key parameters that influence the target speed ranging from slow to fast walking (0.48 and 1.71 m/s) as well as a large range of step lengths (0.43 and 0.88 m) and step duration (0.51, 0.98 s). The findings show that specific reflexes during stance have a major effect on step length regulation mainly given by the contribution of positive force feedback on the ankle plantarflexors’ group. On the other hand, stretch reflexes active during swing of iliopsoas and gluteus maximus regulate all the gait characteristics under analysis. Additionally, the results show that the stretch reflex of the hamstring’s group during landing phase is responsible for modulating the step length and step duration. Additional validation studies in simulations demonstrated that the identified reflexes are sufficient to modulate gait in human locomotion. Thus, this study provides an overview of the possible reflexes to control the gait characteristics.

**Author summary:** 

## 1 Introduction

The neuromusculoskeletal system allows humans and animals to move and interact in their environment choosing among different motor patterns through a complex and redundant interaction of neural circuits. However, the strategies used to control the different gait patterns have not been elucidated yet. It is well-known that the central nervous system controls locomotion in a hierarchical and distributed way by modulating the activity of its control subsystems such as spinal reflexes and central pattern generators (CPGs) ([1], [2]). These networks are modulated by descending cortical and brainstem pathways and sensory feedback in order to regulate the motor outputs for the required motion ([3], [4], [5], [6], [7]). It is also well-established that the modulation of sensory feedback relies on the control of reflex responses through reciprocal and presynaptic inhibition ([8], [9], [10]). In mammals and lower vertebrates, stereotyped movements are executed with low sensorimotor gains, whereas during demanding tasks or unfamiliar conditions locomotion relies on higher feedback gains ([9]). This led to the conclusion that the central nervous system modulates sensory transmission by adjusting reflex gains in order to develop adaptive responses to the environment. Yet, experiments on subjects with lost limb proprioception demonstrated that the amount of motor control delegated to sensory feedback is more prominent in humans compared to other mammals and lower vertebrates ([11]). Therefore, feedback pathways might be more important than central circuits in controlling human biped locomotion. Yet, the way how descending pathways facilitate or inhibit selected spinal circuits in order to modulate gait is still to be fully uncovered.

Neuromusculoskeletal simulations are a powerful tool to test hypotheses in neuroscience and the interaction between biomechanical properties, sensory inputs and spinal circuits ([12], [13], [14]). Several proposed models aimed to reproduce the healthy behavior of human locomotion and its modulation. In chronological order, [15] demonstrated that a simple musculoskeletal model driven by a CPGs network of Matsuoka oscillators ([16]) combined with joint angles and velocity feedback is able to reproduce stable locomotion robust against perturbations. This model is able to change speed by modulating the tonic input to the CPGs and decreasing step duration and stride length. More recently, a reduced neural control strategy has been proposed for the CPG structure based on muscle synergies ([17], [18]). [19] showed that this control scheme generates walking and running patterns at different speeds with the modulation of seven key control parameters. These studies demonstrated how the modulation of selected feedforward components in the spinal cord can reproduce a wide range of walking behaviors. However, these models do not take into account the potential effect of reflex circuits’ modulation in changing the gait characteristics.

On the other hand, other studies highlighted the contribution of sensory feedback to motor control. One of the first contributions in this direction was given by [20] with a neural controller composed of motorneurons receiving inputs from a common CPG and reflexes from stretch and force receptors. Additionally, in this model the spindle reflexes included inhibitory inputs to antagonist muscles and the parameters were optimized using genetic algorithm optimization. Successively, [21] developed a purely reflex-based neuromechanical model encoding principles of legged mechanics reproducing kinematics, dynamics and muscle activation of human walking behavior without the contribution of any CPG circuit. The modulation of speed for this model was explored by [22]. Performing different optimizations for 6 different speeds ranging from 0.8 to 1.8 m/s, Song identified 9 key reflex control parameters that show a significant trend together with the increasing of speed. Three key parameters were related to trunk balance, three to stance behavior and three to swing generation. The model could generate speed transition from slow to fast speed optimizing the identified parameters. However, these key parameters do not include only reflex mechanisms but also balance, prevention of overextension and reciprocal inhibition mechanisms that are simplified in a way that can hardly be related to specific physiological proprioceptive information.

Then, [23] added CPG models to Geyer’s reflex controller to test the hypothesis that CPGs could simplify the control of speed. The CPG models were based on abtract oscillators that could replicate the steady-state muscle stimulation patterns generated by the reflexes during a gait cycle. Therefore, the optimization of reflex parameters was performed once in order to generate stable locomotion and learn the stimulation patterns. Then, the feedback parameters were kept constant and the modulation of the gait was controlled only by tuning the amplitude and the frequency of the oscillators. In particular, it was shown that it is possible to modulate the gait speed by regulating the oscillatory parameters related to the hip muscles. Subsequently, [24] developed a similar controller to generate movement in a simulated biped robot. The CPG model was composed of Matsuoka oscillators and played a major role in controlling the stimulation of proximal muscles, whereas the activation of distal muscles relied on positive force feedback. Also in this case, key parameters for gait modulation were identified performing several optimizations at various speeds. The ones that showed a more relevant trend according with changing of gait speed were the stimulation gains controlling the amplitude of oscillators located at the hip muscles ([25]). Indeed, previous animal experiments ([26]) led to the hypothesis that there exists a proximo-distal gradient in joint neuromechanical control where hip and knee joints are primarily controlled by feedforward circuits whereas distal joints rely more on sensory feedback, especially load sensors. The aforementioned studies investigated the shared control of CPGs and reflexes. Yet, the changing of gait characteristics relies mostly of CPGs parameters and the contribution reflex modulation has not been investigated.

Subsequently, [27] added supraspinal layers on top of a generalized reflex model in the three dimensional space ([28]) able to reproduce walking and running behaviors. These behaviors were achieved through the optimization of stance reflex parameters and the modulation of two supraspinal parameters: desired foot placements and the desired minimum swing leg length. Therefore, the modulation of gait relies on the integration of these descending pathways without identifying and selecting the feedback circuits that contribute to this modulation. Moreover, some components of the control mechanisms are still hard to translate in physiological meaning. More recently, [29] developed a more detailed reflex-based controller modeling each control components basing on physiology of proprioception with the only non-physiological components taking care of trunk balance. The model is designed to walk in the sagittal plane and it is optimized for different target speeds between 0.50 m/s and 2.00 m/s reproducing experimentally observed kinematic, kinetic, and metabolic trends of human walking. This present study will use the model proposed by Ong in order to identify the reflexes taking part in the modulation of speed, step length and step duration.

Indeed, the aforementioned studies demonstrate that both feedback and feedforward controllers are able to faithfully reproduce various walking behaviors highlighting the complexity of the neural and musculoskeletal systems as highly redundant mechanisms ([30], [31]). CPG-based models partially uncovered the contribution of spinal feedforward oscillatory mechanisms generating diverse walking behaviors with the modulation of CPGs parameters. However, previous reflex-based model could not identify physiologically relevant reflex parameters responsible for gait modulation. Furthermore, previous studies in neuromechanical simulations focused mainly on achieving different target speeds rather than separate the components controlling step length and step duration. The relations between these gait characteristics have been investigated by [32] in an experimental study where subjects walked with combinations of slow, nominal and fast step lengths (i.e. 0.584, 0.730 and 0.876 m) and large, nominal and small step duration (i.e. 0.65, 0.52 and 0.43 s). The resulting range of speed is between 0.89 and 2.04 m/s. The study showed that the activity of gluteus maximus, gluteus medius, vastus, gastrocnemious and soleus are the muscles dedicated to vertical support and forward progression independently from changing on step length or step duration and that increased step length results majorly from the larger contribution of hip and knee extensors.

This study aims to understand the potential mechanisms of reflex modulation behind various behaviors of human locomotion. Other mechanisms such as modulation of CPG circuits and muscle synergies could surely play an important role in modulating locomotion, but the present goal is to focus on the potential role of reflex modulation in gait adaptation. In particular, the main focus is dedicated to the modulation of speed together with the independent modulation of step length and step duration. In the present study, we aim at answering the following questions:

1. To which extent can the modulation of reflexes modify speed, step length and step duration during walking?
2. Can these quantities be controlled independently?
3. Which specific reflexes should be modulated to adjust each quantity?

This study will use neuromechanical simulations of human walking performing three different sets of optimization having various target speed, step length and step duration from the lower to the upper boundaries of a human model driven by the reflex controller proposed by Ong. The results suggest that the reflex-based model can generate different gait behaviors including low and high values of speed, step length and step duration. Furthermore, walking patterns ranging among small and large step lengths could be achieved maintaining the step duration fixed and vice versa. Finally, all these behavior can be controlled with the modulation of 9 identified reflex parameters that showed the highest correlation with the changing of gait characteristics.

## 2 Materials and methods

The analysis performed in this study is conducted using the optimization and control framework SCONE ([33]). The musculoskeletal model and the reflex controller are based on the ones used by [29] and are described in more detail in the following sections starting with the presentation of the musculoskeletal model, the reflex controller, the optimization protocol, the description of dataset analysis and the validation steps.

### 2.1 Musculoskeletal model

The musculoskeletal model (Fig 1) is based on the one developed by [34] composed of a skeleton of height = 1.8 m and weight = 75.16 kg. The model movement is constrained in the sagittal plane and has 3 degrees of freedom (DoFs) at the pelvis and other 3 for each leg: one at the hip, one at the knee and one at the ankle. Three spheres are also included as contact model in order to estimate the ground reaction forces when they are in contact with the ground. The contact model is taken from [35] and is composed of one bigger sphere of radius equal to 5 cm at the anatomical reference of calcaneus and two smaller of radius 2.5 cm at the anatomical reference of toes. The model is also composed of nine Hill-type muscle tendon units ([36]) per leg: gluteus maximus (GMAX), biarticular hamstrings (HAMS), iliopsoas (ILPSO), rectus femoris (RF), vasti (VAS), biceps femoris short head (BFSH), gastrocnemius (GAS), soleus (SOL), and tibialis anterior (TA).

**Fig 1.**
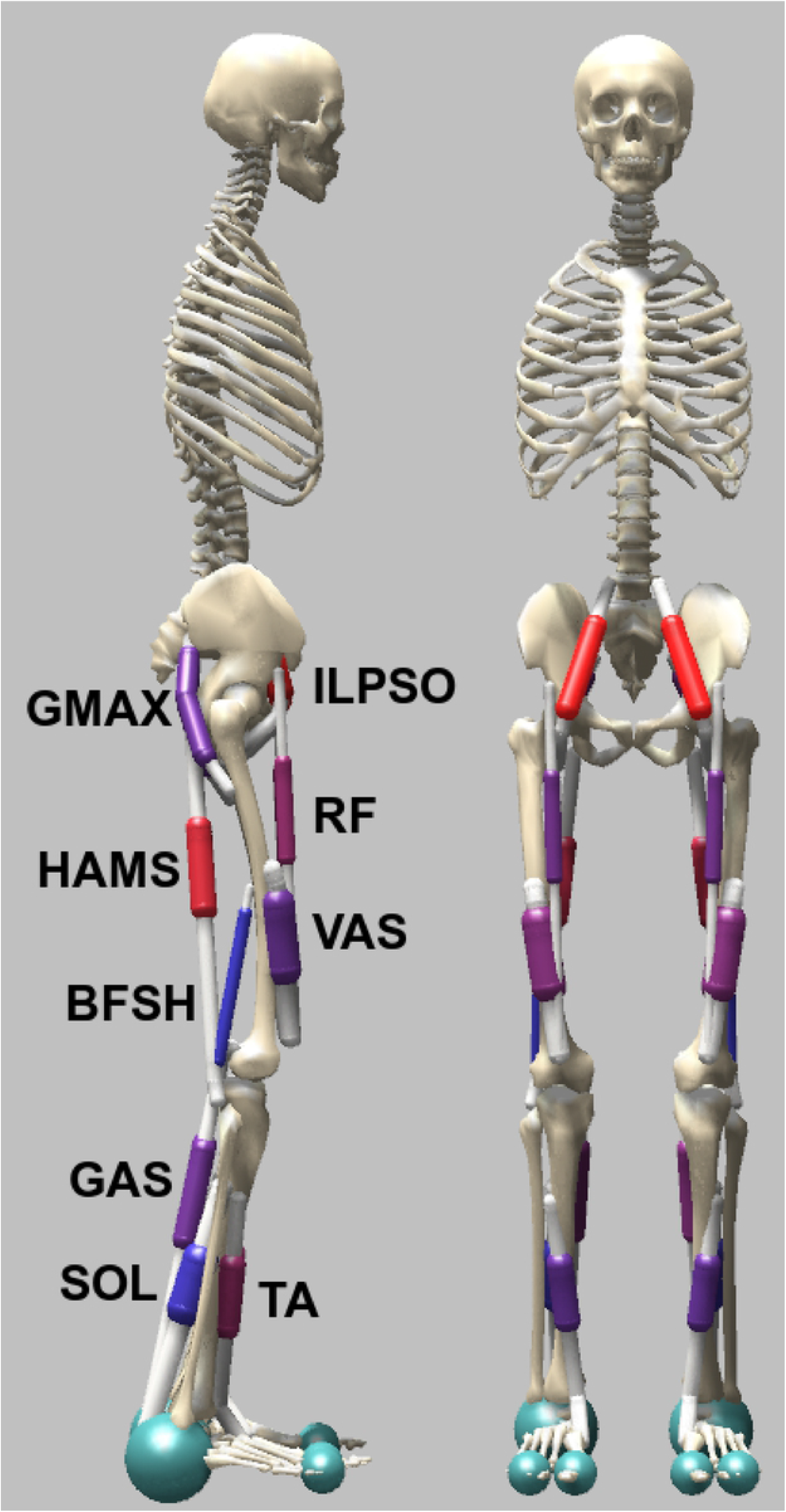
Musculoskeletal model used to study human locomotion. The model is constrained in the sagittal plane and has 9 DoFs: hip and knee flexion/extension, ankle plantar/dorsal flexion for each leg and 3 additional DoFs located at the pelvis:. The movements are generated throught the activation of 9 muscles per leg: gluteus maximus (GMAX), biarticular hamstrings (HAMS), iliopsoas (ILPSO), rectus femoris (RF), vasti (VAS), biceps femoris short head (BFSH), gastrocnemius medialis (GAS), soleus (SOL), and tibialis anterior (TA).

### 2.2 Reflex controller

In the reflex controller proposed by [29], the type of stimulation provided to each muscle depends on the phases of the gait cycle. The gait cycle is divided in 5 different gait subphases, 3 for the stance phase and 2 for the swing: early stance (ES), mid-stance (MS), pre-swing (PS), swing (S) and landing preparation (LP). Taking as reference the division of the gait cycle defined in clinical gait analysis ([37]), it is possible to classify the division in subphases proposed by [29] as follow:

- early stance (ES): first double support and early stance in single support
- mid-stance (MS): mid and late stance in single support
- pre-swing (PS): second double support
- swing (S): early and middle swing
- landing preparation (LP): late swing

The controller is based on three different kind of feedbacks: positive force feedback from the Golgi tendon organs’ Ib fibers, and stretch reflexes length and velocity feedbacks from the muscle spindles’ Ia fibers. Furthermore, PD controllers regulating the forward lean angle of the trunk are integrated in the stimulation of the hip muscles to maintain balance. A constant feedforward stimulation only dependent on the state of the gait cycle is also integrated. The types of stimulation provided to muscles are mathematically described in the equations below:

Feedforward stimulation:

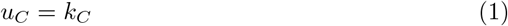

Ia length feedback:

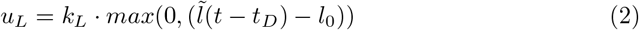

Ia velocity feedback:

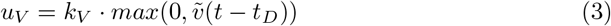

Ib force feedback:

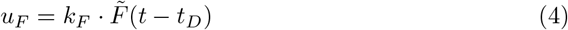

PD balance controller:

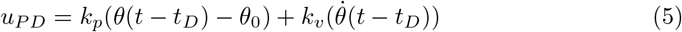

where *k_C_, k_L_, k_V_, k_F_, k_p_*, and *k_v_* are the gains of the reflex controller, *l*_0_ is the length offset of the stretch response and *θ*_0_ is the proportional feedback of *θ*. On the other hand, *t_D_* represents the parameter for the time delay and it depends on the muscle proximity to the vertebral column: *t_D_* = 5*ms* for the hip, *t_D_* = 10*ms* for the knee, and *t_D_* = 20*ms* for the ankle. The variables used in the controller (muscle length *l*, contraction velocity *v* and force generated *F*) are taken normalized according to specific muscle parameters: optimal length (*l_opt_*) and maximum isometric force (*F_max_*).

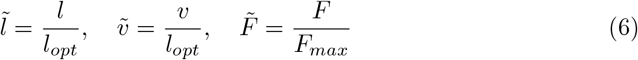

Finally, a state controller regulating thresholds parameters that define the switching between one sub-phase of gait and another is also integrated.

### 2.3 Optimization

The Covariance Matrix Adaptation Evolutionary Strategy (CMA-ES) method is used to optimized the parameters and obtain the different walking behaviors. In each simulation the model walks for 15 seconds unless the model falls. The falling condition is detected recording the height of the model and the simulation stops if its value decays below 0.8 m. The parameters of the optimization are the maximum number of generations equal to 1500, the samples per iteration λ = 16 and the step size *σ* = 1.

Three different sets of optimizations with separate additional objectives in the fitness function are launched. These optimizations are performed with different targets implemented in the objective functions:

- Optimization set 1: different target speeds ranging from slow to fast gait
- Optimization set 2: different target step lengths ranging from small to large maintaining a fixed value of step duration
- Optimization set 3: different target step duration ranging from small to large maintaining a fixed value of step length

The first set of optimizations minimizes the difference between the average speed of the model and a defined target speed. The target speed changes in every optimization covering a wide range, from the slowest to the fastest speed that the stability of the model can handle. The following optimization for step length and step duration are performed starting from the initial condition of the best solution found in the mid-range at 1.0 m/s of speed with a value of step length and step duration around 0.7 m and 0.7 s, respectively. Therefore, the second set investigates the modulation of step length having this gait characteristic as target varying in the different optimizations and a fixed target step period of 0.7 s that does not change among the optimizations. Similarly, for the third set, the step length is kept fixed to 0.7 m and the step duration is the varying target. Both the fixed target step length and step duration are considered with tolerance factors of 0.02 m and 0.02 s. Since forcing a fixed step length or step duration while varying the gait characteristics increases the effort of the task, the weight for effort minimization is reduced from 1 to 0.1 for the last two sets of optimizations in order not to penalize the task’s achievement. The different target objectives for the three sets of optimization are reported in Table 1.

**Table 1.** Different target objectives for the 3 optimization sets: the first raw describes the target ranges that the model is able to bare maintaining stability, the second raw indicates the chosen weight for effort minimization lower for high demanding tasks and the third and fourth raw indicates the additional target of fixed step duration and step length, respectively. The allowed ranges of step length and step duration for set 1 and set 2 are very small in order to modulate one gait characteristic minimizing the changing of the other.

|  | set 1 | set 2 | set 3 |
| --- | --- | --- | --- |
| Target ranges ($w = 100$) | [0.45, 1.75] [m/s] | [0.45, 0.95] [m] | [0.50, 1.00] [s] |
| Effort minimization weight | 1.0 | 0.1 | 0.1 |
| Target step duration | none | [0.68, 0.72] [s] | none |
| Target step length | none | none | [0.68, 0.72] [m] |

On the other hand, the objectives in the fitness function in common with all the optimizations are the avoiding of falling recording the height of the model at the end of the simulations, minimization of overcoming of joints limits and the head stabilization maintaining the vertical and horizontal acceleration of the head in a defined range. These conditions were already implemented in [29] and are reported in Tab 2.

**Table 2.** Common objectives among the 3 optimization sets: the first column indicates the different objectives, the second the corresponding target value and the third the weight assigned in the cost function to the specific objective. The common objectives included a measure to avoid falling solutiocorrere ai riparins (termination height), the minimization of passive torques and the head stability to avoid excessive acceleration of the head

|  | value | weight (w) |
| --- | --- | --- |
| Termination height | 0.8 [m] | 100 |
| Target passive torques | 0.0 [$N \cdot m$] | 0.1 |
| Head stability-y | [-4.9, 4.9] [$m/s^2$] | 0.25 |
| Head stability-x | [-2.45, 2.45] [$m/s^2$] | 0.25 |

### 2.4 Dataset analysis

The three set of optimizations obtained contains several solutions where the achieved values of speed, step length and step duration are recorded together with the amount of effort and passive torques. Specifically, set 1 contains 147 solutions extracted from 12 optimizations, whereas set 2 and set 3 respectively contain 134 and 75 solutions both extracted from 9 optimizations from each set. These selected solutions satisfy the following conditions

- stability: from the solutions that reached the maximum simulation time without falling condition, the stable solutions are the ones that show a convergence toward a constant oscillation of joint angles.
- cost of transport: the efficient solutions from energy point of view are the ones with a cost of transport lower than 4 *J/(kg · m)* for speed between 0.8 and 1.3 m/s and lower than 8 *J/(kg · m)* for slower and faster speeds since these types of gaits requires a higher energy expenditure
- joints limits: the solution extracted should not overcome the joint ranges. The model assigns a penalty depending on how much and how frequently a joint range is overcome. The solutions selected for the analysis are the ones that have a penalty lower than 0.5 for the knee and hip angles while the penalty limit for the ankle angle is set to 3 since the model tend to have a high dorsiflexion.

Additionally, the second and third set of optimization need also to maintain respectively the values of step duration and step length in the ranges reported in Table 1. From the data obtained, it is possible to evaluate the ranges of the three gait characteristics when the movement of the human model is driven by reflexes allowing to answer to the first research question.

For the identification of key parameters, the focus is purely on reflex circuits. Therefore, the parameters related to balance, feedforward stimulation and thresholds are not analysed. The identification of important parameters for gait modulation is found through the analysis of the correlation coefficient between the parameter and the variation of the gait characteristics analysed. Therefore, a reflex parameter is considered a key parameter for gait modulation if it presents a correlation coefficient larger than 0.6 for at least one of the three gait characteristics’ modulation. Then, each identified parameter is analysed through three different regressions, one for the solutions obtained by each of the three optimization sets. The regression analysis is performed with a similar methodology as previously done by [25]. However, we take in consideration all the good solutions extracted instead of their average. In order to evaluate the tendency of data distribution, the data are regressed finding the lowest order polynomial function that is able to model the distribution with a coefficient of determination (*R*^2^) larger than 0.7. In case the solutions extracted are widely spread, the maximum polynomial order allowed is set to 3.

### 2.5 Validation of gait behaviors

The previous stage allowed to identify the parameters that mostly affect gait modulation. However, in case of stretch reflexes the same level of stimulation can be achieved modulating differently the gain (*k_L_*) or the length offset (*l*_0_). Therefore, when one of these stretch parameters is identified as key parameter, the other belonging to the same stretch reflex is also taken into consideration as key modulator for the validation study. Once the key reflexes are identified, we demonstrate that the variation of these is sufficient to achieve the same ranges of gait variability achieved in the previous experiments. This process is done performing new optimizations exploring the boundaries of the gait characteristics (minimum and maximum speed, step length and step duration). During these optimizations, reflexes that were not identified as relevant are not allowed to change and their value is kept constant. On the other hand, the key reflexes are optimized together with the parameters regulating balance, feedforward stimulation and states. A further validation is done optimizing only the non-relevant reflexes for the gait modulation and keeping the key parameters to a constant value with the target objective of obtaining the same boundaries of the gait characteristics obtained previously. This process is done in order to demonstrate the reduced ability of the model in modulating the gait without the possibility to change the key reflex parameters. The other parameters not belonging to the reflex controller are optimized also in this case.

The identification of key reflexes and the validation process permit to answer to the last two research questions allowing to define which reflex controls which gait characteristics and if these characteristics can be controlled independently. Then, the study presents the gait analysis of joints kinematics, ground reaction forces and muscular activity taking in consideration minimum, intermediate and maximum values of the obtained speed, step length and step duration.

## 3 Results

This section presents first the gait limits that the model is able to reach in term of speed, step length and step duration. Then, the identified key parameters are presented. These parameters are divided depending on whether they control step length, step duration or both. The identification of the key parameters is done analysing the linear correlation for the three gait characteristics and the tendency of the data distribution shown together with the regression model. The data used, figures and videos can be found at the following link: https://drive.switch.ch/index.php/s/A34EeDAHhs2xzQR

Three different sets of optimizations were performed. Table 3 shows the minimum and maximum boundaries reached during the optimization processes. From the first one investigating the speed modulation, the solutions were extracted within the range of speed from 0.45 to 1.71 m/s, step length from 0.45 and 0.87 m and step period from 0.51 to 1.04 s. The solutions contained in the second set of optimizations were able to achieve a minimum step length of 0.45 and a maximum of 0.88 m, covering the same range already obtained in the first set with an increasing target speed. The step duration was maintained constant at 0.69 s and this condition has been satisfied for all the solutions selected with a tolerance of 0.01 s. Consequently, the range of speed obtained in this second set is reduced compare to the first one because of the imposed fixed step duration and it is included between 0.69 and 1.48 m/s. Finally, the third set presents solutions with values of step duration ranging from 0.51 to 0.91 s. The target step length is maintained fixed to 0.72 m and the solutions extracted satisfied this condition with a tolerance of 0.02 m. The range of speed obtained out of this optimization set is included between 0.78 and 1.46 m/s.

**Table 3.** Boundaries of the three gait characteristics targeted during the three optimization sets. The first set shows the results for the optimization where the gait target to reach is the desired speed that varied from the lower to the upper boundary. The second set shows the results for the optimizations with a fixed step duration and a varying target step length. Finally, the third set shows the results for the optimizations with a fixed value of step length changing the target step duration.

| Optimization set [min, max] | Speed | Step length | Step duration |
| --- | --- | --- | --- |
| Set 1 | [0.45, 1.71] [m/s] | [0.45, 0.87] [m] | [0.51, 1.04] [s] |
| Set 2 | [0.69, 1.48] [m/s] | [0.45, 0.88] [m] | [0.68, 0.70] [s] |
| Set 3 | [0.78, 1.46] [m/s] | [0.70, 0.74] [m] | [0.51, 0.91] [s] |

The changing of parameters’ values in accordance with the modulation of gait characteristics permits to identify the reflexes controlling speed, step length and step duration. Table 4 presents the correlation coefficients of the 9 key parameters identified Among these, some reflexes have been found to be linked with specific gait characteristics. Specifically:

- 4 key reflexes for the modulation of speed through the only modulation of step length
- 3 key reflexes for the modulation of speed through the modulation of both step length and step duration.
- 2 additional key reflexes modulating both step length and step duration accordingly resulting in a small effect in speed changing
- no reflex parameter have been found to modulate step duration independently from step length.

**Table 4.**
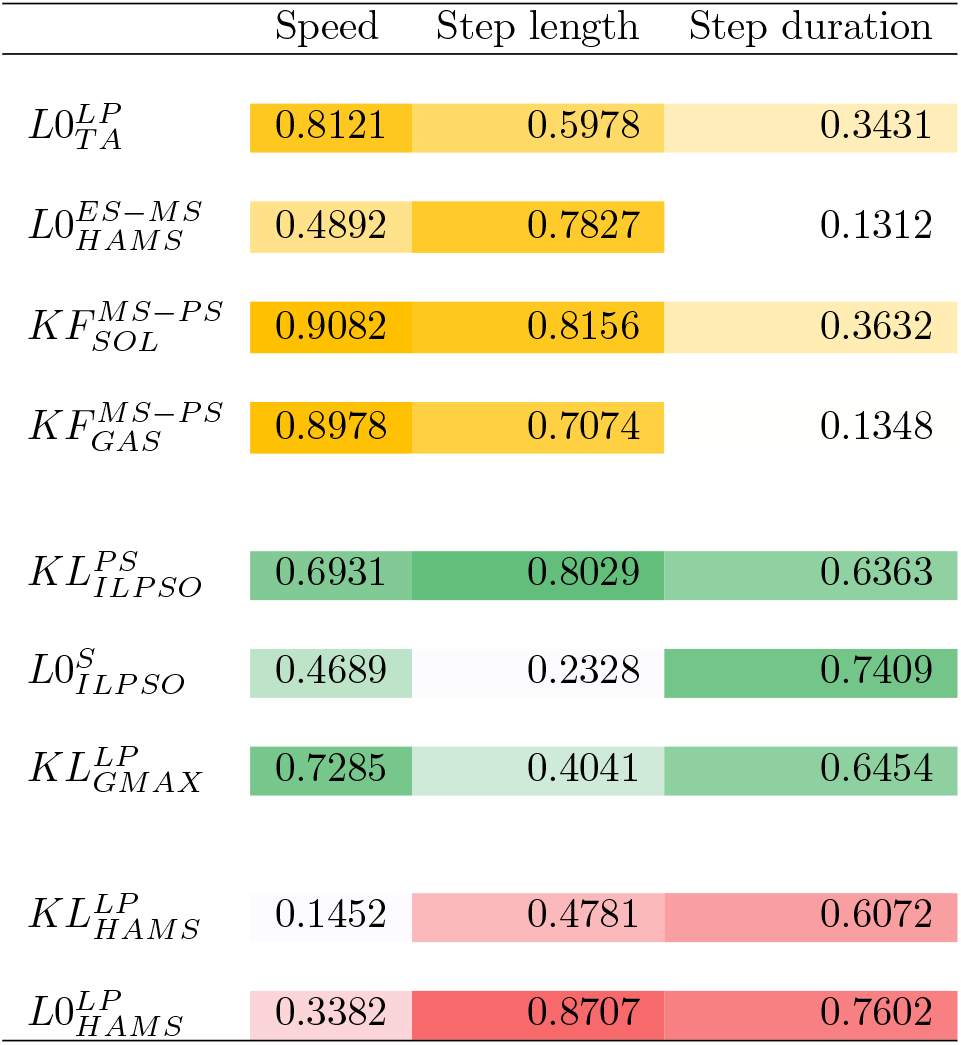
Correlation coefficients of identified key reflex parameters: the 4 reflexes having a major effect on speed modulation through the modulation of step length are highlighted in yellow, the 3 ones having effect on all the three gait characteristics are highlighted in green and the ones modulating step length and step duration accordingly with small effects on speed are highlighted in red.

Since we did not found parameters that where able to modulate step duration without having a significant effect on step length, step duration is mainly modulated by reflexes that also affect step length. Therefore, the solutions presented where the modulation of step duration is achieved maintaining the step length fixed could be obtained only with compensation mechanisms performed by other reflex circuits.

### 3.1 Step length modulators

In total, four parameters have been selected since those are the ones that showed a high correlation coefficient with speed and step length and a low correlation coefficient with step duration indicating a minor effect on this latter gait characteristic. The length offset of tibialis anterior during landing preparation is the only parameter active during the swing phase that has a high correlation coefficient with both speed and step length (c = 0.8121 and c = 0.5978, respectively). From the graphs at the top in Fig 2, it is possible to notice that the speed decreases linearly with the increasing of stretch length offset of tibialis anterior. This tendency is mainly due to coherent decreasing of step length whereas the values of the parameter remains roughly constant for a large range of step durations. Indeed, these values show a significant changing only for very short duration values below 0.55 s proper of fastest gaits.

**Fig 2.**
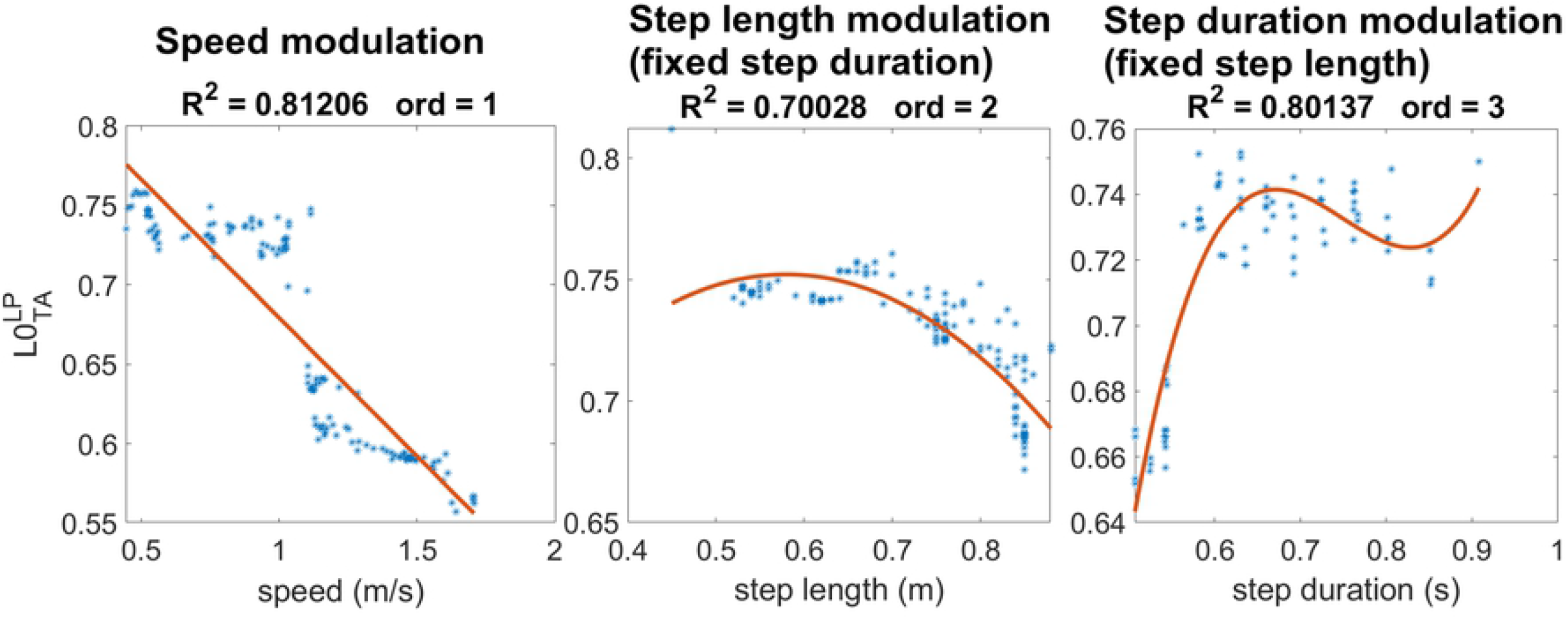
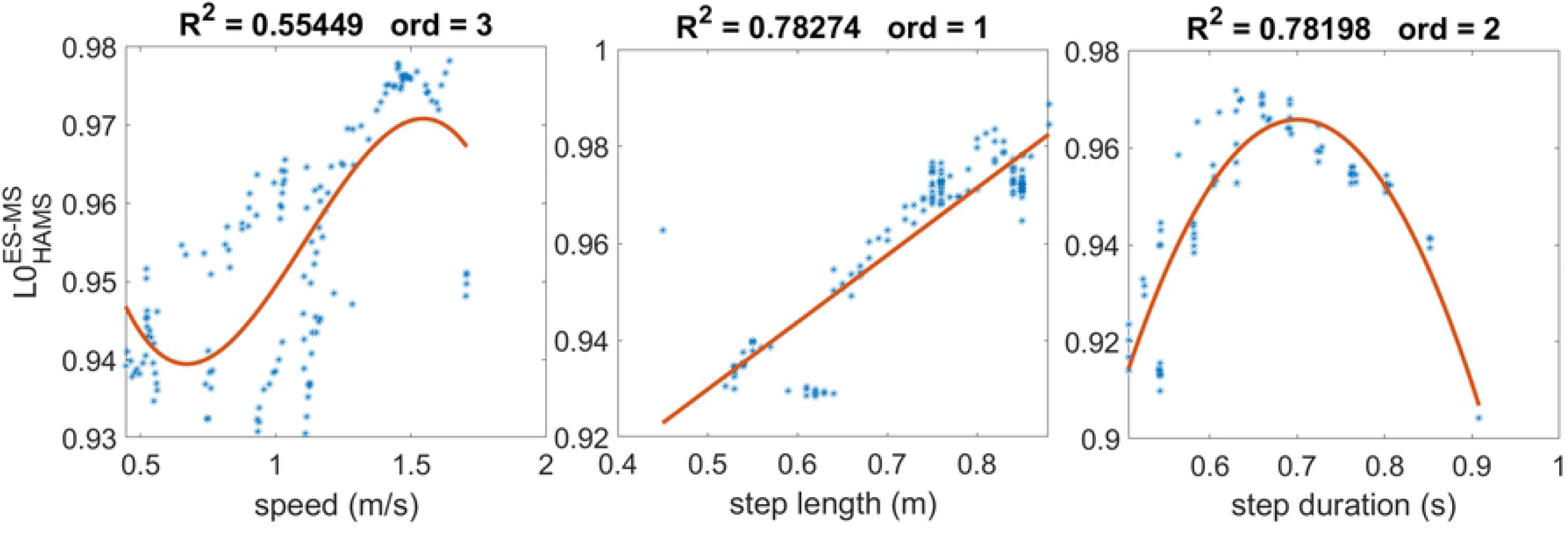
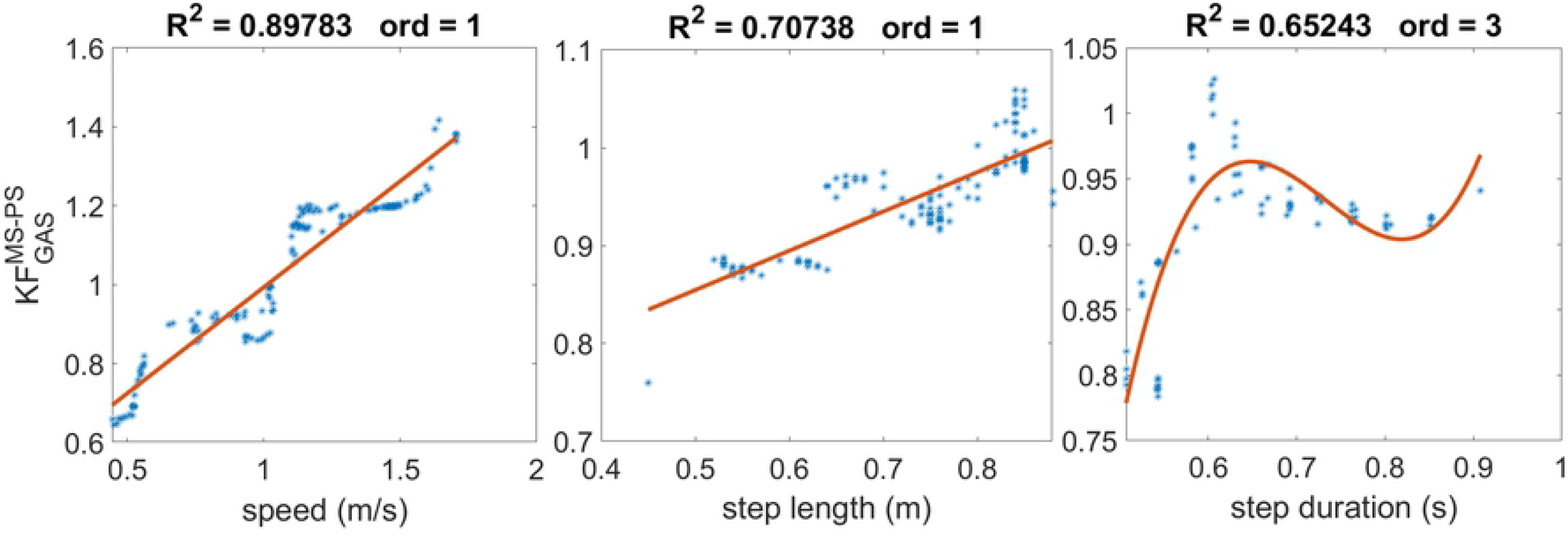
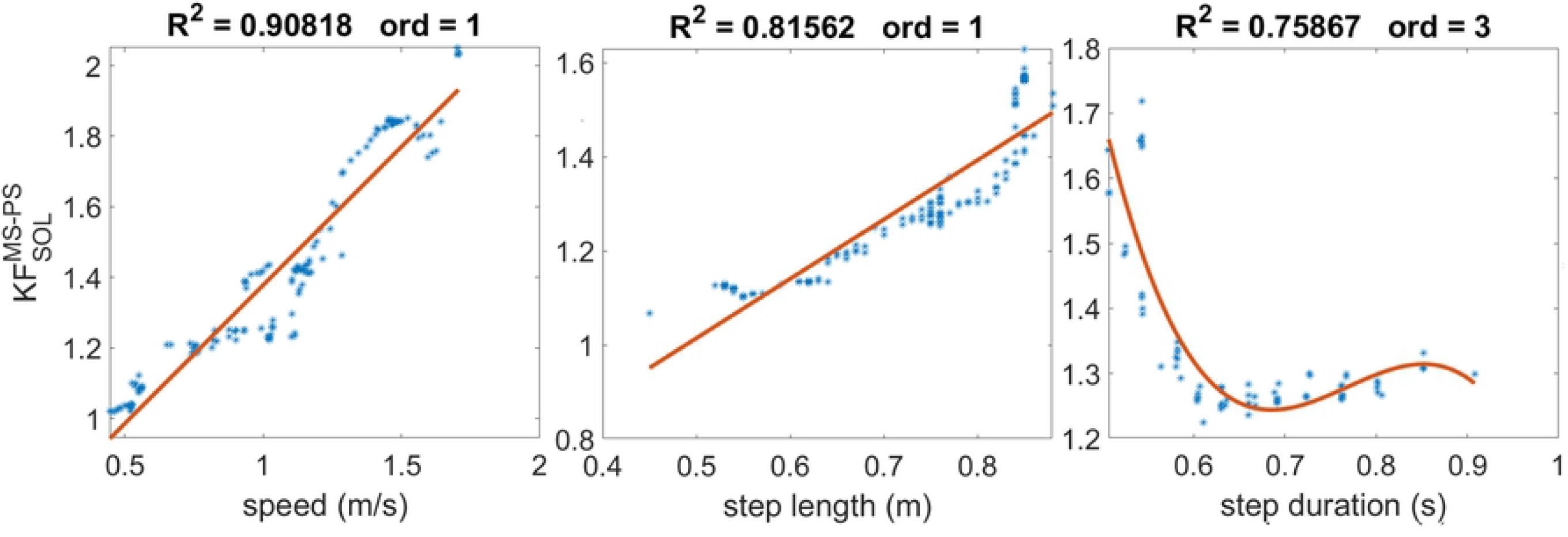
Regression analysis of step length modulators. The solution obtained by the three sets of optimizations are represented by the blue dots, whereas the regression is represented by the red curve. The plot on the left show the data distribution and regression for the speed modulation while the step length and step duration modulation are shown by the plots on the center and on the right, respectively. Length offset of tibialis anterior is the only reflex active in swing that affect step length and speed with a decreasing effect when the parameter’s value increases. By contrast the other reflexes presented facilitate speed and step length increasing with minimal effect on the step period.

Another important reflex parameter is the length offset of the hamstring muscle in the first two sub-phases of stance. The graphs in the second raw in Fig 2 shows the linear relationship with the step length modulation. However, the step duration is also affected by the parameter’s variation as shown by the quadratic regression for its modulation (*R*^2^ = 0.78198) with the peak at the middle of the range. Therefore, the same parameter value can be used to generate two different gaits with different step durations depending on the compensatory mechanisms of the other reflex parameters. This nonlinear effect on the step duration results in a less clear dependency between the hamstrings stretch offset and the increasing of speed. In fact, the values of this parameter are very spread around the linear regression for the speed modulation. However, the global increasing tendency seems to follow the one found for step length modulation.

On the other hand, the propulsive muscles seem to have a key role in the modulation of step length. Indeed, the positive force feedback in stance of both gastrocnemious and soleus have a high linear correlation with step length with *R*^2^ = 0.81562 for soleus and *R*^2^ = 0.70738 for gastrocnemious. This linear dependency can be found also in the modulation of step length that can be modeled as a linear regression with *R*^2^ = 0. 90818 for soleus and *R*^2^ = 0.89783 for gastrocnemious. The changing of these two parameters show to have minor effects on the modulation of step duration with low correlation coefficients and the parameters’ values maintained roughly constant for longer step durations than 0.6 s. Yet, shorter durations more proper of high speeds may have an effect on the parameters’ values. Indeed, the positive force feedback gain of soleus increases when step duration goes below 0.6 s while the one of gastrocnemious decreases.

### 3.2 Step length and step duration modulators

The next results present the key parameters affecting significantly both step length and step duration. These parameters are separated in two different group: the ones that affect speed modulation and the ones that have no effect on speed. This diversification is made because there are possible parameters that influence step length and duration coherently maintaining the ratio between these two gait characteristics roughly constant minimizing the effect on speed modulation.

#### 3.2.1 With effects on speed

The three parameters represented in green in Table 4 are the key reflex parameters that influence both step length and step duration with significant effect on speed. The length feedback of iliopsoas is one of the key modulators. Indeed, the graphs at the top of Fig 3 described the behavior of the stretch reflex gain of the iliopsoas during pre-swing. This parameter decreases linearly with the increasing of step length and increases with the increasing of step duration resulting in a global decreasing of speed.

**Fig 3.**
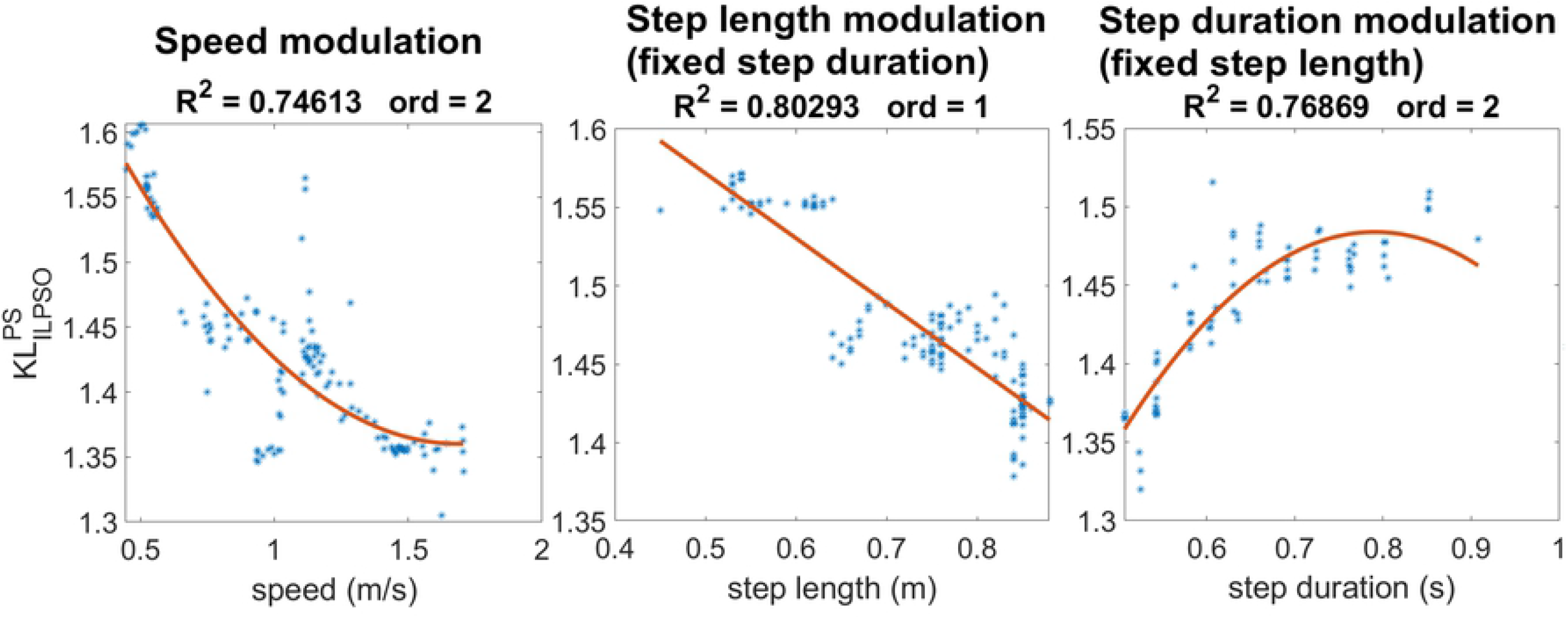
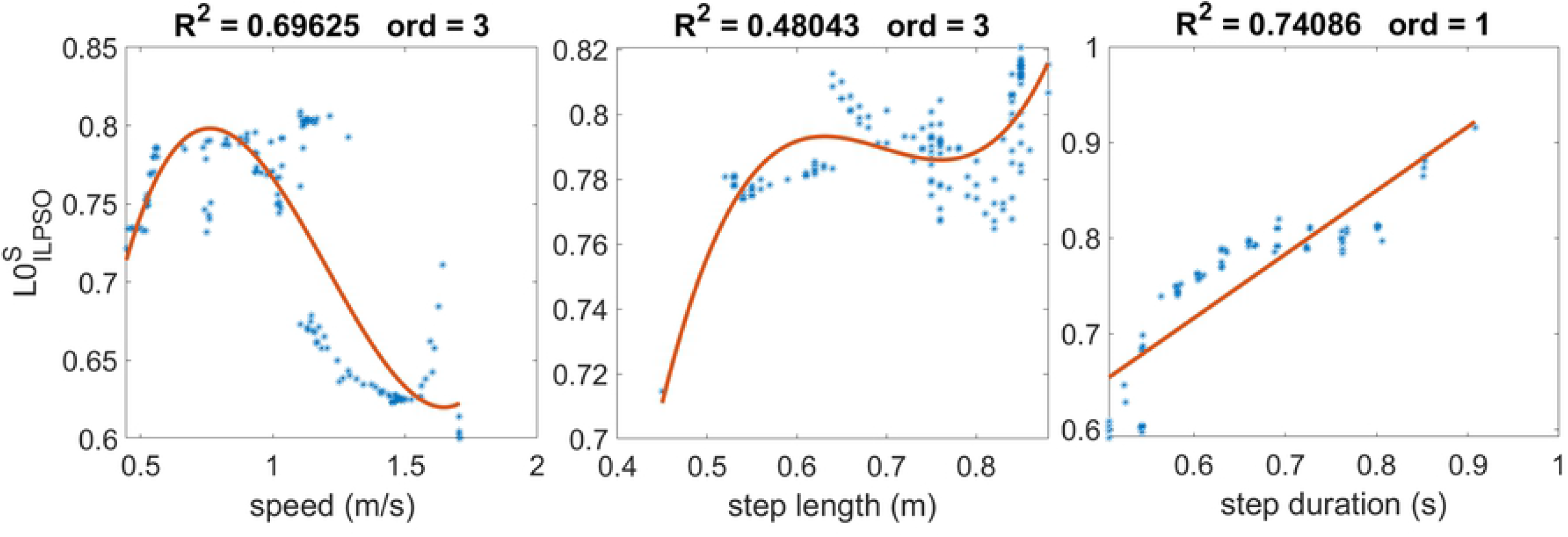
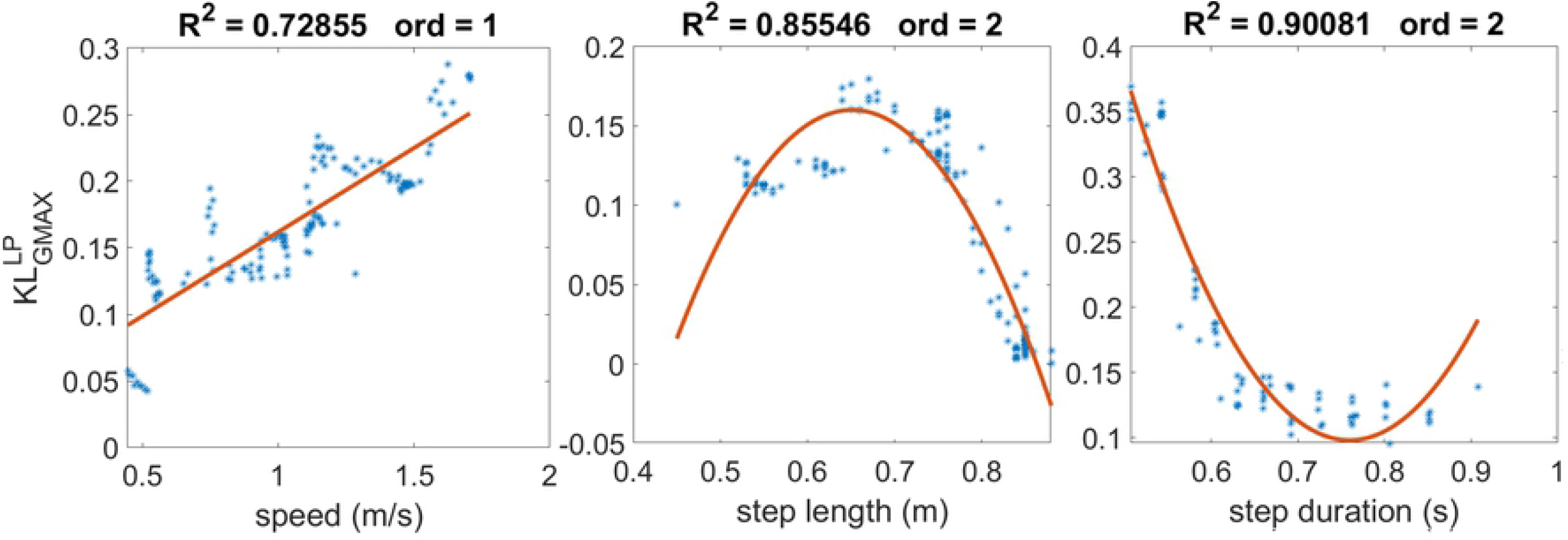
Regression analysis of step length and step duration modulators with effects on speed. The stretch reflex of iliopsoas during pre-swing and the length offset during swing have both a decreasing effect on speed due to decreasing of step length and increasing of step duration for the former and primarily for step duration increasing for the latter. On the other hand, the stretch reflex of gluteus maximus in landing preparation has an linear increasing effect on speed with non-linear effects on step length and step duration

In the following phase of the gait cycle, we can observe a decreasing of speed with the increasing of length offset of iliopsoas during swing, as described in the plots presented in the second raw of Fig 3. The speed decreasing is coherent with the observed tendency of the step duration that increases linearly with the increasing of length offset value. The step length also seems to increase accordingly with the increasing of parameter’s value. However, this increasing is significant only at the extremes of the step length boundaries for step lengths shorter than 0.6 m and longer 0.8 m.

The last parameter presented in the length feedback gain of the gluteus maximus during landing. We can observe the influence of this reflex stimulation on speed that is efficiently modeled by the regression with a linear relationship. This linearity is not found in the modulation of step length and step duration. The important contribution on the modulation of step length is given mainly a step lengths larger than 0.7 m showing a drastic decreasing behavior and a minor increasing for smaller step lengths. This increasing behavior for small step lengths is probably the main contributor of the increasing behavior of speed. However, for fastest speed the values of the parameter keep on increasing despite they decrease for large step lengths. Therefore, at fastest speed the modulation is given by the effect on the step duration that decreases rapidly when the reflex gain increases.

#### 3.2.2 Without effects on speed

Finally, the last two key reflex parameters presented in Table 4 control the modulation on both step length and step duration maintaining their relation roughly constant resulting in a minimal effect on speed. Both these parameters are related to the stretch reflex activity of the hamstring during landing preparation and are represented by the gain and the length offset. From the graphs at the top of Fig 4, the gain decreases accordingly with both step length and step duration resulting in a null effect on the speed modulation as shown by the spread distribution of data and from the low coefficient of determination of the third order polynomial (*R*^2^ = 0.19406). The increasing of length offset values with the increasing of step length and step duration also contributes to the reduces activity of the stretch response when the two gait characteristics increase as shown in the bottom graphs in Fig 4F. Also in this case, the global effect is a less efficiency in the modulation of gait velocity.

**Fig 4.**
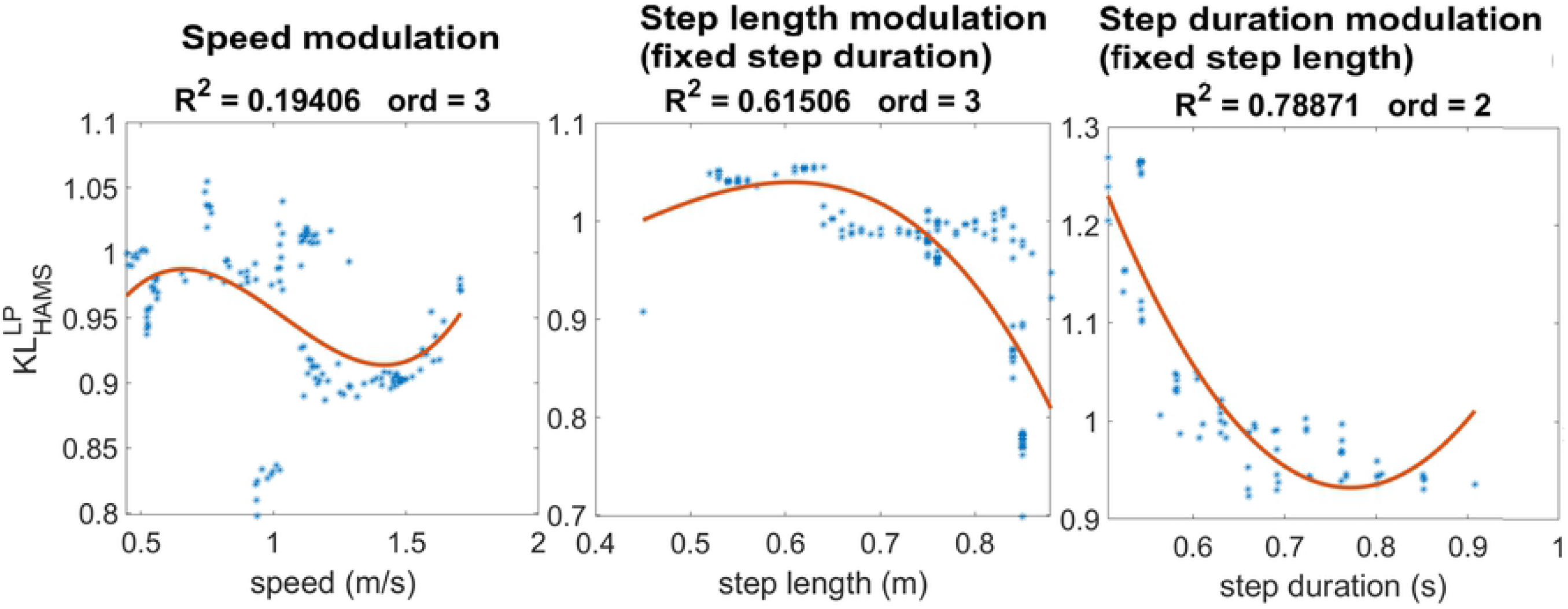
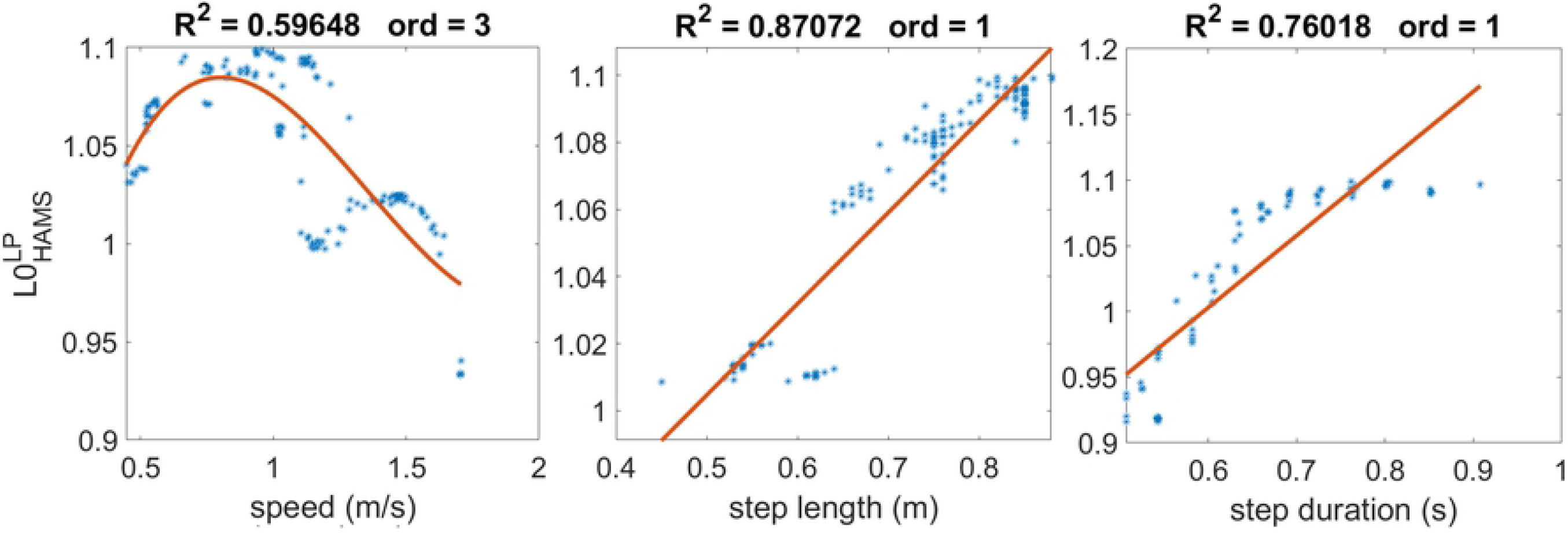
Regression analysis of step length and step duration modulators with effects on speed. Increasing the stretch reflex gain of hamstring during landing preparation leads to a decreasing of step length and step duration resulting in a low influence in speed modulation. Similarly the increasing length offset of hamstring during landing preparation results in an increades step length and step duration with a small effect on speed modulation.

### 3.3 Modulation of key parameters

From the previous section we were able to identify the reflexes that affect mostly the gait modulation. These key reflexes are highlighthed in the control diagram of Fig 5. Step length modulators are highlighted in yellow, whereas step length and step duration modulators are highlighted in green and red depending on whether they have or do not have an effect on speed, respectively.

**Fig 5.**
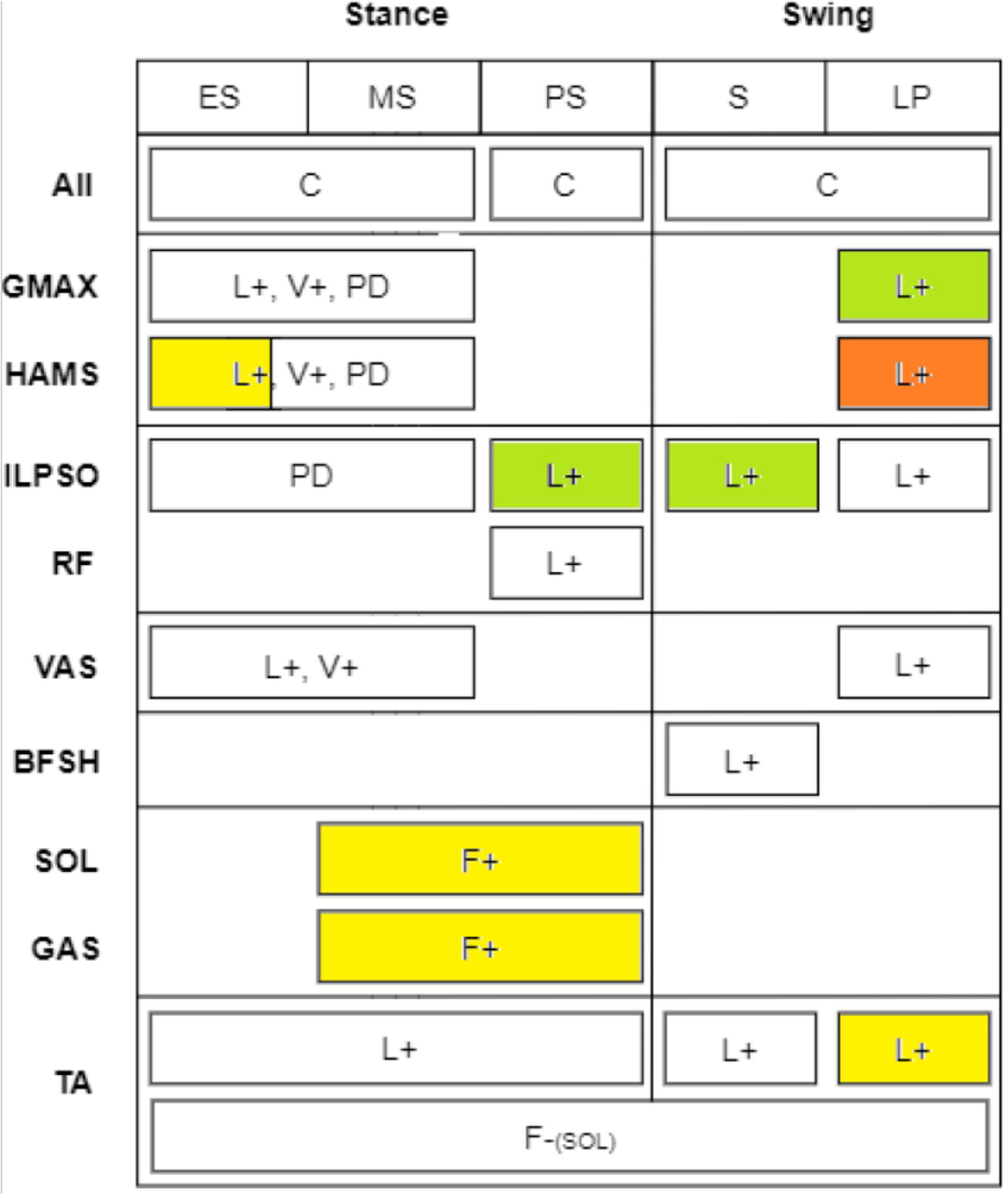
Diagram of reflex controller with key reflexes modulating gait highlighted. The reflexes that were found to modulate mainly step length are highlighted in yellow, whereas the ones that were found to modulate step length and step duration together are highlighted in green and red depending on whether they showed a significant effect on speed (green) or not (red).

Then, in this section we presents the largest ranges reached for the modulation of the three gait characteristics optimizing only the key reflexes identified together with the feedforward, balance and the state controller parameters. These boundaries are compared with the ones obtained previously optimizing all the parameters and with the ones obtained optimizing all the parameters except the key reflexes that are kept constant. Table 5 presents the largest boundaries obtained optimizing the key reflexes identified. Comparing to Table 4, the modulation of key parameters could generate locomotion behaviors from slow to fast gaits with large and small step length and short and long step duration. The boundaries of the three gait characteristics cover the same ranges of the ones obtained optimizing all the reflex parameters. Similar results are obtained for the modulation of step length and step duration. Therefore, the key reflexes selected demonstrated to be able to modulate gait with the same performances of the modulation of all reflexes.

**Table 5.** Boundaries of the three gait measures reached with the optimization of key reflexes. The optimization of the key reflexes alone could obtain the same performances of the gaits obtained optimizing all the reflexes suggesting that the major role of modulation is delegated to the key reflexes.

| Optimization set [min, max] | Speed | Step length | Step duration |
| --- | --- | --- | --- |
| Speed modulation | [0.48, 1.71] [m/s] | [0.43, 0.88] [m] | [0.51, 0.98] [s] |
| Step length modulation | [0.77, 1.26] [m/s] | [0.52, 0.87] [m] | [0.69, 0.70] [s] |
| Step duration modulation | [0.79, 1.30] [m/s] | [0.70, 0.71] [m] | [0.54, 0.91] [s] |

However, some reflexes that were not considered as key modulator could still have significant effects on the modulation of locomotion since the neural system is highly redundant. The results from the optimization of non-relevant reflexes targeting the same ranges achieved in the previous stages show that the model fails to achieve slow and fast speed targets when the key identified reflexes are kept constant and not included in the optimization. For step length modulation maintaining a fixed value of step duration, the model is not able to reproduce stable locomotion with small step length. However, large step gaits could be achieved with similar performances obtained previously also without the modulation of the identified reflex modulators. On the other hand, the optimization could not reach long step durations maintaining to an intermediate value of 0.67 s when trying to target high values. Yet, the optimization could converge to a behavior able to reproduce short durations comparable to the ones previously obtained. These results suggest that there are parameters beyond the key reflexes identified that can modulate large step length and short step duration. These parameters do not necessarily belong to the reflex controller, but could be part of the feedforward, balance or state controller. In order to verify this, we performed optimizations varying the key reflex parameters alone and other optimizations varying only the other reflexes maintaining constant in all cases feedforward, balance and state controller parameters. We verified that the modulation of state controller parameters alone is able to achieve large step lengths. By fixing these parameters we could reach a step length value of 0.87 m with the optimization of key parameters, whereas the optimization of other reflexes could only reach a step length below 0.8 m. On the other hand, the modulation of short step duration could be achieved with the contribution of balance parameters. Maintaining these parameters constant, short step duration of 0.54 s could be achieved optimizing the key reflex parameters while the other reflexes could not converge to solutions with a step duration value lower than 0.62 s. Therefore, the key reflexes identified describes a large variance of the modulation of the neural feedback mechanism, whereas the other reflexes do not seem to affect significantly gait modulation.

### 3.4 Gait analysis

This section briefly describes the results of the gait analysis changing the three gait characteristics. Additional figures and more details are provided in the Supplementary Information. It can be firstly noted that the reflex controller could generate human-like locomotion behaviors as shown in Fig 6 for the specific solution from the first set of optimization at the intermediate speed of 1.2 m/s.

**Fig 6.**
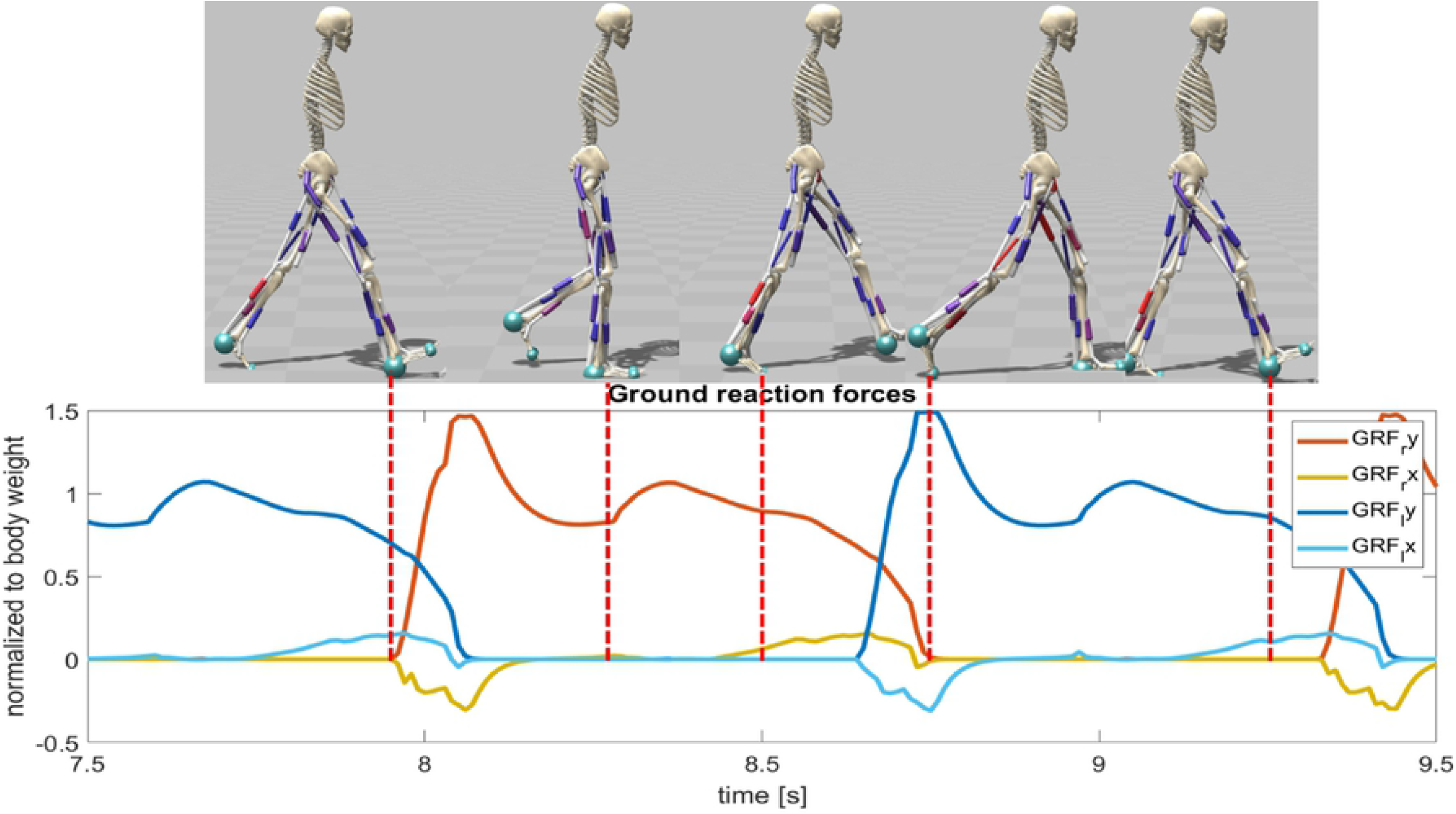
Representation of gait behavior with snapshots taken at different frames of for the speed modulation solution with target speed equal to 1.2 m/s. The instant taken is represented together with the behavior of ground reaction forces where the right foot is the reference and red dash lines correlate the instants of GRFs with the right foot position in the simulation.

The modulation of step length appears to influence more the changing of joint angles oscillations more for hip and ankle angles rather than knee flexion. Furthermore, from ground reaction forces, it can be observed an increased time ratio between stance and swing phases when the model walks at slow speed and small steps, whereas the increasing of step duration does not influence this ratio. Some limitations are found in the excessive dorsiflexion of the ankle over all the gait cycle and the shape of ground reaction forces that presents high peaks for fast speeds and long steps and a second peak slightly anticipated in the gait cycle.

From the analysis of muscle activation, we can observe that the muscle mostly affected by the modulation of the gait are the hamstrings, the iliopsoas and the gastrocnemious having an increased activity with the increasing of gait speed. In addition, some muscles in which there was no changing of reflex parameters because not considered relevant, could still present an increased activity for faster walking patterns. An example of this can be found in the activation of vasti that exhibits large variations depending on the target gait despite its reflexes being kept constant and not modulated.

More details and figures can be found in the Supplementary information section

## 4 Discussion

In this study we aim first to understand how much a human model controlled by sensory-driven neural signals alone is capable to replicate various behaviors of gait at different speed, step length and step duration. The results obtained from the optimizations show that large ranges of these three gait characteristics could be generated by the reflex controller. These results are coherent with the previous studies involving the application of sensory-driven controllers ([21], [27], [29]) focused mainly on the single modulation of energy efficient walking at different speeds. Our results demonstrated that sensory reflexes are able to modulate not only speed and energy efficient gaits but could also to control step length and step duration independently generating behavior beyond the optimal energy efficiency that humans are able to perform even if these patterns do not represent the best strategy.

Then, the last research question aims to investigate if specific reflexes could present a major effect in modulating specific gait characteristics. Nine key reflex parameters are identified, 4 of which modulating mainly step length, 3 modulating all the three gait characteristics and 2 modulating step length and step duration accordingly with small effects on speed. These results demonstrate that the modulation of the key reflexes is sufficient to generate various behaviors of human locomotion ranging from reduced to high values of speed, step length and step duration similar to the ones obtained with the optimization of all the reflexes. Therefore, the modulation of a small subset of reflexes could possibly be involved in the strategies used by descending commands to change the gait behavior together with the already investigated modulation of feedforward circuits ([23], [19]).

Some of the identified parameters active mainly during stance were found to have a primary effect on the step length. Indeed, step length is affected by the level of propulsion that the muscles of the stance leg can give pushing the body forward and lowering the center of mass. Coherently with experiments in human subjects ([38]), the main propulsion is given by the soleus and gastrocnemious muscles through their positive force feedback as it can be observed by the strong correlation that the two reflexes have with both speed and step length modulation. However, the muscle activation of soleus and gastrocnemious show a considerable changing only with the modulation of speed and no meaningful variations are observed with the modulation of the step length. It should be observed that the muscle activation depends not only on the values of reflex parameters, but also on the state of the muscle itself as observed for the activation of the vasti not dependent on the variation of reflexes.

It has been verified that the positive force feedback gains of soleus and gastrocnemious change accordingly with the data distribution and regression laws described in the results section. Therefore, the unchanged level of activation observed for the modulation of step length is due to an alteration of muscle states. Another parameter affecting the step length is the length offset of the hamstrings during the stance phase. More precisely, an increase of the hamstrings length offset results in a faster gait with larger steps and a decrease in a slower gait with small steps. Indeed, the length offset defines the level of length for the muscle fiber after which the stretch reflex is active. Therefore, a larger length offset allows the hamstrings to sustain a level of stretch due to the knee extension and hip flexion happening majorly in early stance at higher step lengths without a large response of the stretch reflex that would produce an undesired knee flexion.

On the other hand, the modulation of gait with significant effects in all the gait characteristics relies largely on reflexes active during swing phase or swing preparation. The main stretch activity that plays a role in this modulation is the one governing the activity of iliopsoas. In fact, the larger length offset of this muscle causes a slower response of the stretch activity resulting in a slower and lower leg lifting typical of gait with reduced speed. Moreover, the decreasing stretch activity of iliopsoas during the swing preparation when the step length increases allows a full hip extension that is necessary for larger steps.

Then, the fast execution of landing is guaranteed with the modulation of gluteus maximus and hamstring stretch reflexes. The activity of the gluteus maximus during the landing phase is important to determine the step period since its higher stretch response allows a faster landing of the foot increasing the frequency of the gait. On the other hand, The stretch reflex of hamstring during landing phase regulates the coherent increasing or decreasing of step length and step period through the regulation of both reflex gain and offset. The increased activity of hamstrings helps a faster landing phase due to the hip, but it also prevents a full extension of the knee reducing slightly the step length. The excessive knee flexion is prevented by the regulation of the length offset that tend to increase linearly with the increasing of step length. Yet, this regulation tend to slow down the stretch response of the reflex that is less effective in a fast landing.

The modulation of reflex parameters described above involve the regulation of sensory-motor gains (*k_F_* and *k_L_*) and the threshold for the onset of the stretch response (*l*_0_). Physiologically, the modulation of gains can be interpreted with the involvement of presynaptic inhibition of afferent activity ([39], [40]). In addition, the regulation of descending modulation and *γ*-motoneurons contribute to the modulation of stretch reflex threshold. The altered regulation of this reflex component takes also a role in the generation of motor impairments in gait pathologies ([41]).

[32] studied the effects of step length and step duration in lower limb muscles. They observed that gluteus maximus, gluteus medius, vastus, gastrocnemious and soleus are the most important muscles for the forward progression. In particular, it was shown that the muscles mainly responsible for the modulation of step length are the hip and knee extensors. In the current study, the musculoskeletal model does not include the gluteus medius, but some of the other muscles identified by the aformentioned study are activated by the key reflex parameters selected for gait modulation and others show different activities despite their reflexes are not modulate such as the vasti. Indeed, the modulation of gait is mainly regulated by gastrocnemious, soleus, hamstrings, gluteus maximus and iliopsoas reflexes. In the results concerning the muscle activation we also observe a major involvement of hip and knee extensors as reported in the experimental study. Looking at the modulation of reflexes, the circuits governing the activity of soleus and gastrocnemious positive force feedbacks also have a high correlation with the changing of gait characteristics, but plantarflexor muscles are considered less important. However, it should be observed that the current study has different ranges of speed, step length and step duration explored. In fact, they did not cover the range of slow walking with the minimal speed recorded at 0.89 m/s and the minimal step length of 0.58 m. On the other hand, our model was able to walk at 0.45 m/s with a short step length of 0.45 m and a large step period of 1.04 s. Therefore, the ankle plantarflexors are probably more important for gait modulation if very slow ranges are also included. It has been verified that the reflex parameters related to hip and knee extensors has significant trends in the modulation of step lengths only for larger steps whereas they remain roughly constant for smaller steps resulting in a low coefficient of determination in the regression study. Furthermore, another experimental study conducted by [42] demonstrated that the activity of distal muscles like soleus and gastrocnemious has much less variability among subjects in slow walking compared to proximal muscles like semitendineous and gluteus maximus. Therefore, slow gait speeds rely mostly on defined activity of propulsive muscles rather than hip muscles.

Other experimental studies investigated the variation in muscles activity and in the activation of spinal cord regions at different speeds. [43] showed that with increasing in speed the muscles activation increases especially for the distal muscles active in stance like gastrocnemious and soleus, whereas the increasing activity of hip muscles is less consistent especially for the iliopsoas where the recorded activity is kept low for every speed analysed. From our results we could replicate the increasing activation of muscle plantarflexors but we also observe a large increase in activation of iliopsoas muscle. It should be noticed that experimental studies using surface EMGs are limited from the muscle deepness. Indeed the iliopsoas is located deep in the trunk and it is difficult to record its activity [44]. In addition, [45] estimated the activity of the motoneurons in the spinal cord following the work by Ivanenko and found that the activation ratio between lumbar and sacral segments is increasing with the increasing of speed. Therefore, proximal muscles controlled by lumbar segments increase their activity more consistently than distal muscles controlled by sacral segments. We also observe this condition in our results where there is a more important increase of iliopsoas and hamstrings muscle activity compared to the increasing activity of soleus and gastrocnemious when passing from slow to fast speeds.

Concerning the kinematics, previous studies investigated the different kinematic behavior when speed is changing from slow to intermediate. More importantly, [46] performed experiments on healthy subject walking from very slow speeds (0.1 m/s) to self selected speed. These experiments showed that the stance phase duration in percentage of the gait cycle duration pass from 70% in slow speeds around 0.5 m/s to 60% in self selected speed as also observed in our study. In addition, the larger oscillations between flexion and extension for the hip angle at higher speeds are also found in the experiments. Similar observations can be done for the ankle angle even though our results present an excessive dorsiflexion compared to the experimental study. For the knee angle, the current study did not found significant changes on the knee flexion in swing among different speed. Similarly, the study found a significant reduction of knee flexion only for very slow walking close to 0.1 m/s. Yet, the controller used in the current study is not able to reproduce very slow gaits below 0.4 m/s where the contribution of feedforward neural mechanisms are probably more important than the feedback mechanisms. In the end, the ground reaction forces obtained and described before have the limitation to present a high initial peak when the foot strikes the ground and the anticipated second peak that are not found in experimental studies. This behavior of ground reaction forces together with the excessive dorsiflexion of the ankle is probably due to local minima found in the optimization. However, we found higher peaks with the increasing of speed with almost flattened peaks for slow solutions as also described in Wu’s study.

Finally, from the validation performed, the variation of the key reflexes achieved the same gait behaviors obtained optimizing all the parameters of the controller. Furthermore, not allowing the key parameters to change resulted in a drastic limitation of gait modulation. Physiologically, it is possible that the neural system modulates additional feedback circuits for human locomotion than the one identified. Yet, the results suggest that the identified reflexes are sufficient to modulate the gait in the contest of purely sensory-driven mechanisms. However, feedforward neural system also plays an important role in the regulation of walking behavior. Abstract models of central pattern generators basing their rhythmic patterns on sensory feedback and muscle synergies have already demonstrated to be able to modulate walking and running behaviors ([23], [24], [19]). Future works should focus on the combined modulation of feedforward and feedback circuits with detailed models of the spinal cord as already implemented for mouse models ([47]) investigating other aspects of modulation of human walking involving higher voluntary control such as high ground clearance, stairs climbing, walking on slopes and obstacle avoidance.

## Conclusion

In this work, we investigated the nature of sensory reflexes modulation behind the generation of different walking behaviors using neuromuscular simulation tools. We focused on the study the modulation of speed, step length and step duration identifying the main reflexes that were found to be responsible for the modulation of the three gait characteristics. Hamstrings length stretch reflexes and plantarflexors positive force feedbacks during stance were found to have a primary effect on step length modulation together with tibialis anterior length stretch reflex during landing. On the other hand, stretch reflexes of iliopsoas, hamstrings and gluteus maximus active during pre-swing, swing and landing were found to modulate both step length and step duration. These reflexes demonstrated to be sufficient and necessary to modulate a wide range of the three gait characteristics under analysis. Furthermore, the solutions obtained showed similarities with previous experimental studies on gait modulation concerning kinematics, ground reaction forces and muscle activation ([32], [43], [45], [42], [46]). This study provides a first contribution of the modulation of human locomotion in simulation environments based on physiologically relevant neural feedback circuits. Future directions should focus on investigating joint contribution of feedforward and feedback neural components in the modulation of human gait.

## Supporting information

### Supporting information related to this publication

Details on gait analysis for low, intermediate and high values of each gait characteristics.

